# EPICURUS: E-field based spatial filtering procedure for an accurate estimation of local EEG activity evoked by Transcranial Magnetic Stimulation

**DOI:** 10.1101/2025.02.16.638512

**Authors:** Xavier Corominas-Teruel, Tuomas P. Mutanen, Carlo Leto, Maria Teresa Colomina, Cécile Gallea, Martina Bracco, Antoni Valero-Cabré

## Abstract

**Background:** The concurrent use of Transcranial magnetic stimulation and electroencephalography (TMS-EEG) is increasingly integrated into research and clinical protocols. However, a reliable isolation of EEG responses that are locally evoked by TMS at the targeted cortical sites independent from contaminating sources, remains challenging.

**Methods:** Here we introduce EPICURUS, a novel spatial filtering approach for TMS-EEG that uses individualized MRI-based simulations of the TMS-induced electric field (E-field) to define the spatial extent of locally evoked activity. This method guides the reconstruction of EEG signals originating from the direct stimulation site while minimizing crosstalk from distant, non-targeted sources.

**Results:** In synthetic simulations and a human TMS-EEG dataset, EPICURUS preserved early-latency TMS-evoked local activity while substantially attenuating later components, consistent with suppression of non-local activity.

**Conclusion:** By leveraging the spatial precision of individualized E-field modeling, EPICURUS may enhance the specificity of EEG signal reconstruction, offering a promising tool for improving the spatiotemporal resolution of local early and late cortical local responses directly elicited by TMS.

## INTRODUCTION

Concurrent Transcranial Magnetic Simulation and Electroencephalography (TMS-EEG) allows for the direct assessment of cortical reactivity and excitability with high temporal resolution [1,2]. However, TMS-evoked potentials (TEPs) often lack spatial specificity, as they reflect a complex mixture of neural sources activated both directly and indirectly by stimulation. Beyond the local cortical response, TMS-evoked activity propagates to regions connected to the stimulated area and engages additional auditory, tactile, and vestibular pathways, activating secondary cortical areas unrelated to the target. Consequently, commonly used measures of cortical excitability, such as TEPs peak amplitudes or latencies, are intermixed with non-local activity, especially at later latencies (> 100 ms) [3–7], limiting the accurate extraction of physiologically meaningful local signals.

To address this limitation, substantial efforts have been made to reduce peripheral contamination, both by taking care of limiting contaminating sources during data acquisition and/or via post-hoc signal processing [5,8–10]. Experimental approaches, such as noise masking or sham procedures, have been proven valuable but often fail to fully suppress multisensory responses or may inadvertently alter TEP profiles. Signal-processing approaches, including sensor-level signal examination, sensor-level spatial filters (e.g., Laplacian filters) [11–13], source separation techniques or source-based artifact-removal strategies [14,15] aim to isolate components of interest within the mixed TMS-evoked response. However, these methods have shown limitations. Sensor-level filters typically employ the same configuration across participants and therefore do not account for individual differences in the stimulated cortical site. In contrast, blind separation approaches, which often rely on arbitrary decisions regarding component selection and rejection, may inadvertently remove genuine TMS-evoked activity [16]. Despite these efforts, disentangling local from non-local TMS-evoked activity (including multisensory responses) without affecting signal integrity remains a largely unresolved challenge. To date, no method has been explicitly optimized to isolate local TMS-evoked activity in the presence of concurrent activity from multiple distant dipoles that may overshadow the local signal.

Here, we introduce **EPICURUS** (**E**-field-based s**P**atial f**I**ltering for **C**oncurrent EEG-TMS **U**sed to ta**R**get so**U**rce **S**ignals), an approach designed to isolate the activity locally elicited by TMS. Our approach defines the targeted cortical region based on participant-specific TMS electric-fields (E-field) distributions while minimizing contributions from distal sources. Specifically, it employs a source-space inverse operator that combines individualized E-field forward simulations [17] with a minimum-norm estimator constrained by crosstalk functions (CTFs) [18–20]. To assess its performance, we employed synthetic data, simulating response to single TMS pulses across different cortical regions, as well as an experimental human TMS-EEG dataset, targeting the primary motor cortex (M1). We hypothesized that EPICURUS would preserve activity originating from a primary stimulated area while suppressing non-local contributions. Specifically, we expected this filter (1) to preserve the amplitudes of early-latency TEP peaks (<100 ms), reflecting local activity; and (2) to attenuate the amplitudes of late-latency TEP peaks (>100 ms), which are typically dominated by distal (e.g., multisensory) contributions.

## RESULTS AND DISCUSSION

To assess the performance of EPICURUS across different target regions, we generated synthetic data that modeled TMS-evoked activity within the left M1 (L-M1), the left dorsolateral prefrontal cortex (L-DLPFC), and the left intraparietal sulcus (L-IPS) [21]. For each target area, we simulated cortical evoked responses at the site of stimulation, mixed with multisensory responses (from the somatosensory and the left and right auditory cortices; see the supplementary Materials for details). For each site, we generated 100 trials, implementing trial-to-trial amplitude variability (random amplitude variance of ±0.5 µV). We then extracted trial-level information from the signal in each target region across all trials using 4 signal extraction strategies: (1) Sensor-level: raw signals extracted from the electrode closest to the stimulated target (hence, C3 for M1, F3 for DLPFC, and P4 for IPS); (2) Laplacian: signals extracted from the same electrode subtracting the average of its immediate neighbors to increase local contrast; (3) Standard MNE: mean source signals (Classical L2-MNE based) extracted from an ROI defined by the TMS E-field magnitude (sources exceeding 80% of the E-field; [18,20]); and (4) EPICURUS: source signals extracted with our E-field-based spatial filtering approach optimized using CTF functions, implementing distributed source constraints to minimize signal leakage across the whole brain and explicit constraints targeting both auditory regions to suppress their contributions.

We compared signals extracted with the four approaches using non-parametric cluster-based statistics (repeated-measures dependent-samples ANOVA [22,23]), analyzing each target area separately. Highly similar patterns of effects across all stimulation targets were found, with significant clusters indicating differences across the four signal-extraction approaches at early (∼20-40 ms), mid (∼60-80 ms), and late (∼130-210 ms) latencies (all p values ≤ 0.03; Figure 1; please see Supplementary Table 2 & 3 for extended results). EPICURUS was the only approach preserving for all targets, the amplitude of early responses while simultaneously suppressing late-latency clusters dominated by signals originating outside the targeted area (e.g., auditory cortex). At early latencies (∼21-45 ms), EPICURUS signals were significantly larger in amplitude than those extracted with Laplacian and Standard MNE approaches (EPICURUS *vs*. Laplacian: *t* = 2.5-6.1; *vs*. Standard MNE: *t* = 2.0-6.5; all *p* values < 0.001) and comparable to single-sensor signals (all *p* values > 0.05). Conversely, at late latencies (∼130-210 ms), EPICURUS more effectively suppressed later peaks compared to all other approaches (EPICURUS *vs*. Standard MNE: *t* = 2.0-5.4; *vs*. Electrode: *t* = 2.6-6.7; *vs*. Laplacian: t = 2.5-3.7; all *p* values ≤ 0.02). Highlighting its ability to isolate local cortical activity with a good signal-to-noise ratio, these findings indicate that EPICURUS was particularly well suited to preserve early local TMS-evoked responses while selectively suppressing late, non-local components.

**Figure 1.**
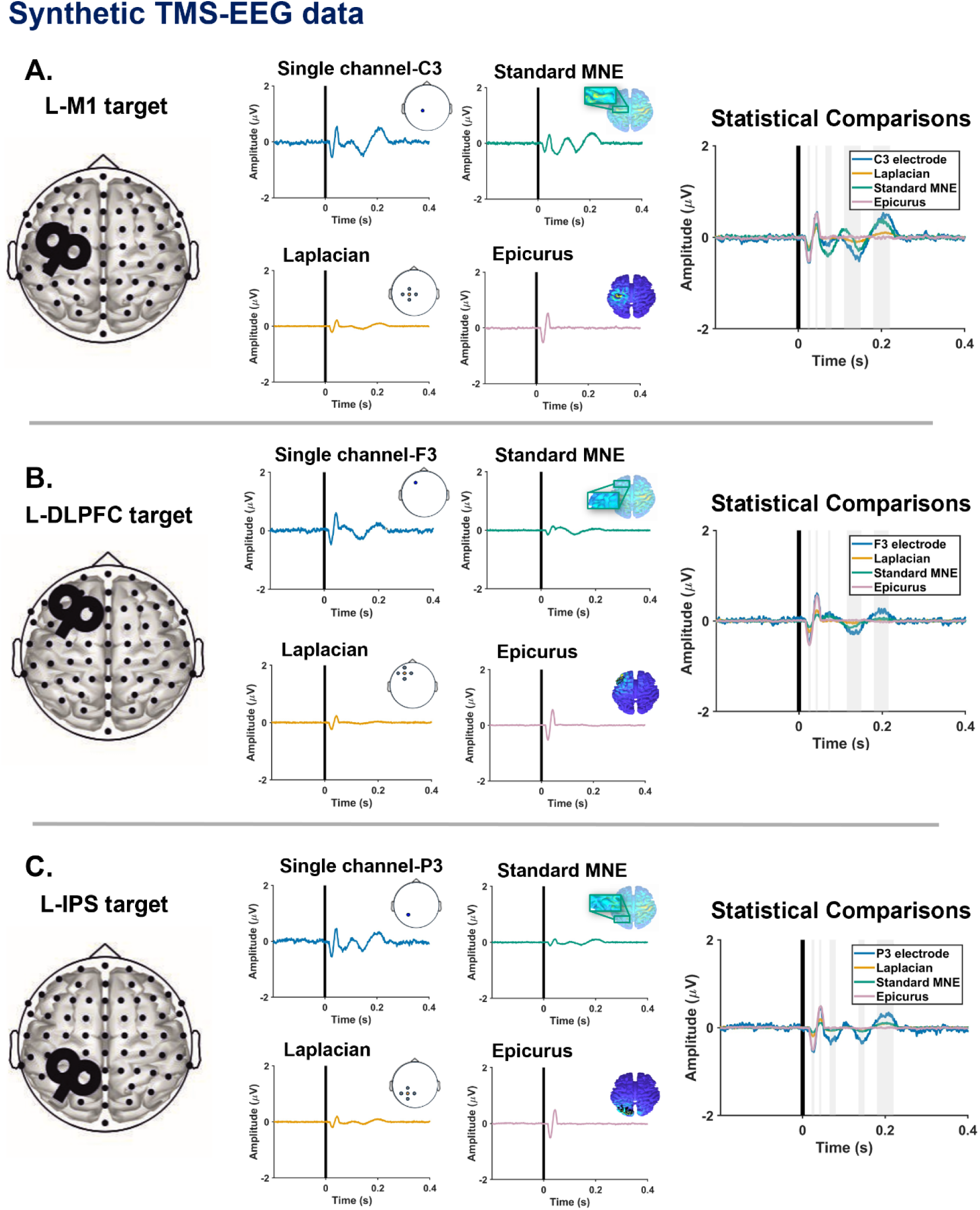
***Synthetic TMS-EEG data.*** Synthetic datasets were generated to benchmark EPICURUS against ground-truth signals across three distinct targets: the left primary motor cortex (L-M1; **A.**), the left dorsolateral prefrontal cortex (L-DLPFC; **B**.), and the left intraparietal sulcus (L-IPS; **C**.). Left column: 3D brains represent the targeted location on each simulated dataset (black coil). Center columns: Time-series for each method are displayed alongside cortical maps showing the spatial filter constraints: single sensor closest to the target (blue), Laplacian (yellow), Standard MNE restricted to the E-field ROI (green), and EPICURUS (pink). Right column: Statistical comparisons highlight significant time-domain clusters (grey shaded bars) where the four methods diverged (p < 0.05, cluster-based repeated-measures dependent-samples ANOVA). Vertical black lines at t = 0 s represent the simulated TMS pulse.

We subsequently evaluated EPICURUS in a pre-existing human TMS-EEG dataset [24] to provide a more realistic and complex test case than synthetic simulations based on isolated cortical dipoles. Eighteen participants (10 females, 8 males, 26±5 years old) received TMS over the left-M1 while EEG was concurrently recorded. Following preprocessing, we obtained TEPs for each participant, using the same four signal-extraction strategies described previously (see the supplementary Materials for full details). Cluster-based permutation statistics using repeated-measures ANOVA [22,23] identified five significant clusters spanning early (∼15-40 ms), mid (∼95-115 ms), and late (∼185-325 ms) latencies (F(3,51) = 4.3-8.5, all p values < 0.02; Figure 2; see Supplementary Table 2 & 3 for extended results). Replicating the pattern observed in synthetic data, EPICURUS effectively preserved early-latency TEPs while suppressing later, spatially diffuse activity. At early latencies (∼15-45 ms), EPICURUS signals were significantly larger than those of the Laplacian and also Standard MNE approaches (EPICURUS vs. Laplacian: t = 2.1-7.6; vs. Standard MNE: t = 3.3; all p values ≤ 0.02) and comparable to single-sensor signals (p > 0.05 at the earliest latencies, except at ∼40 ms in which EPICURUS slightly reduced the amplitude, t = −1.2, p = 0.02). Conversely, at mid-to-late latencies (∼110-325 ms), EPICURUS demonstrated robust signal suppression compared to both the electrode and Standard MNE (EPICURUS vs. Electrode: t = 8.7-15.2; vs. Standard MNE: t = 4.3-7.7; all p values < 0.001).

**Figure 2.**
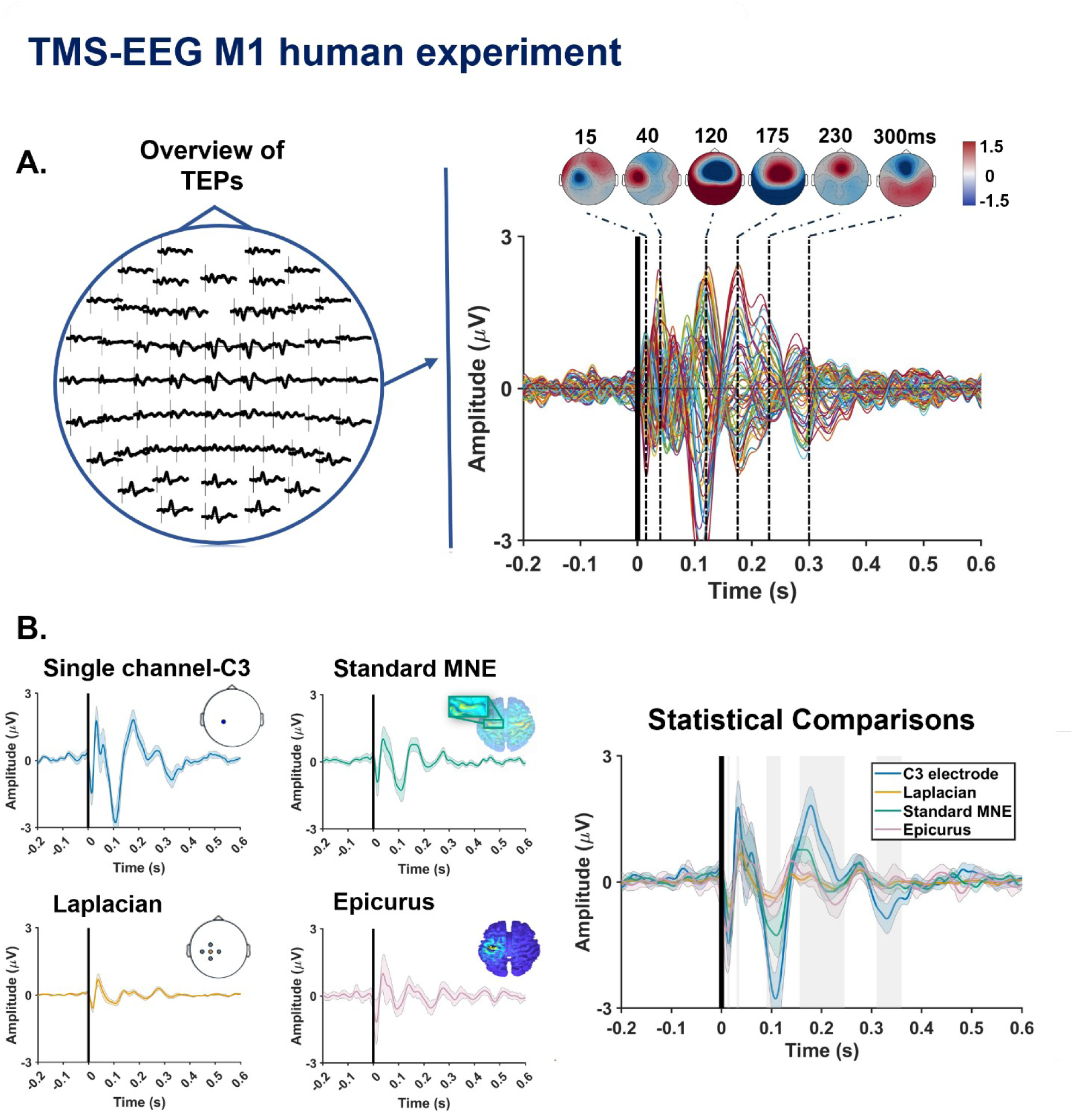
***TMS-EEG M1 human data*.** TEPs were first characterized at the sensor level. **A**; The circular insert on the left displays the electrode-level TEP waveforms across the scalp. On the right, an overview of the TEPs is shown via a butterfly plot of all channels and their associated scalp topographies at key latencies (15, 40, 120, 175, 230 and 300 ms). **B**; Bottom Left: Time-series extraction using the four tested methods: Single channel C3 (blue), Laplacian (yellow), Standard MNE restricted to the E-field ROI (green), and EPICURUS (pink). Small inserts for Single channel C3 and Laplacian indicate the sensor-level selection, while the cortical maps for EPICURUS and Standard MNE visualize the spatial filter distribution and the E-field-informed ROI, respectively. Gray-shaded areas around the traces represent the standard error of the mean (SEM). Bottom Right: Statistical comparisons illustrate significant time-domain clusters (grey shaded bars) where the signal extraction methods diverged (p < 0.05, cluster-based repeated-measures ANOVA). Vertical black lines denote the TMS pulse at t = 0 s.

Overall, our results strongly suggest that EPICURUS preserved early-latency TEP amplitudes (< 100 ms) while selectively attenuating later components in both synthetic and human TMS-EEG data. Performance was particularly robust in in-silico simulated data, as in the human M1 ones, early positive MEP peaks (∼40 ms) were partially attenuated compared to single-channel electrodes. This likely reflects the inherent difference in complexity between the simulated and real TMS-EEG data. While the first of these responses originated in synthetic data solely from the targeted area, in the human brain real TMS evokes activity in adjacent regions via thalamo-cortical loops, near-instant cortico-cortical propagation, and electric field spread. Consequently, EPICURUS may have treated some of these ‘near-neighbor’ physiological signals as off-target activity, leading to its partial suppression. Despite, EPICURUS still significantly outperformed standard MNE and Laplacian approaches, demonstrating its efficacy in preserving early-latency TEPs.

Regarding late TEPs, EPICURUS successfully and consistently suppressed late components in both datasets. In the human data, this was especially notable as late-latency activity consists of a complex mixture of indirect multisensory responses and direct, long-range neural propagation to areas anatomically or functionally connected to M1. This suppression was achieved through two complementary mechanisms: by attenuating signals according to their distance from the target, the filter reduced direct propagation to distal areas; additionally, whereas by targeting specific regions such as the auditory cortices, EPICURUS masked known sources of indirect multisensory activation. In this regard, it is important to note that a key technical constraint arises when TMS is applied near cortical regions that elicit strong peripheral sensory responses (e.g., the somatosensory cortex). In these instances, EPICURUS will recover neural activity from the target region but will not be able to distinguish between the brain’s direct response to the magnetic pulse and indirect multisensory responses if they overlap. Because these types of responses would overlap spatially within the stimulated site, EPICURUS would likely fail to suppress somatosensory responses. For such contexts, Blind Source Separation (BSS) approaches combined with rigorous experimental controls may be more appropriate [7,11].

Ultimately, compared to sensor-level, Laplacian, or standard MNE approaches, EPICURUS provided a superior balance between preserving early local signals and suppressing distal contributions. This makes this filter particularly valuable for studies focused on local responses, for which traditional source-space methods may underestimate early activity due to over-regularization or spatial leakage. Researchers interested in cortico-cortical connectivity or long-range propagation may extend the EPICURUS framework by integrating anatomical spatial priors (e.g., fMRI, diffusion imaging) or applying multiple simultaneous spatial filters to distinct connected regions.

In terms of practical applications, computing the source-space inverse operator is relatively time efficient (∼3 min), though it requires the prior generation of individualized head models (e.g., *Charm* routine in SimNIBS [25]) and E-field simulations, which can take 1-4 hours depending on available processing power. However, once the spatial filter has been derived, its application to EEG data has a rather negligible computational cost, similar to conventional sensor-level Laplacian filtering (e.g., Hjorth filtering). Additionally, compared to Laplacian filtering, EPICURUS offers distinct advantages for subject-specific customization and the ability to enforce custom CTF constraints. The combination of modest computational costs and spatial specificity makes EPICURUS suitable for a wide range of paradigms, including real-time, EEG-informed TMS applications.

In conclusion, EPICURUS appears to be a spatial filtering approach optimized to suppress non-local brain activity while preserving physiologically meaningful local signals. Despite further validation in real biological data from across different brain regions may prove helpful, the method has displayed, according to our analyses, robust performance in both simulated and real human M1 recordings, hence showing promise for the characterization of the early and late local oscillatory dynamics that underlie TMS-evoked brain activity.

## DATA AND CODE AVAILABILITY

The datasets produced and/or analyzed during the current study cannot be made publicly available due to institutional regulations. However, they can be obtained from the corresponding authors of the manuscript upon reasonable request. MATLAB codes for the pre-processing and filtering implementation can be found at the permanent link: https://github.com/XavierCorominas/EPICURUS_E_field_eeg_spatial_filtering

## CREDIT AUTHORSHIP CONTRIBUTION STATEMENT

Conceptualization: XC-T, TM, MB, AV-C. Methodology: XC-T, TM, MB. Data curation and analysis: XC-T. Funding acquisition: CG, MB, AV-C. Supervision: MB, AV-C. Writing - original draft: XC-T, AV-C & MB. Writing - review and editing: XC-T, TM, CL, CG, MT-C, MB, AV-C.

## ACKNOWLEDGEMENTS

This work was supported by the Agence Nationale de la Recherche in France (BrainMAG; ANR-19-CE37-0021; AV-C), the European Union’s Horizon 2020 research and innovation program under the Marie Skłodowska-Curie grant agreement 897941 (MB), the Big Brain Theory (BBT-3; FORTE) internal ICM grant (MB, CG & AV-C), ICM CARNOT training grant (MB & CG), IHU-ICM CARNOT Maturation grant, and additional funding came from Investissements d’avenir (ANR-10- IAIHU-0006) awarded to XC-T and AV-C for associated projects.

## SUPPLEMENTARY MATERIALS

### METHODS

#### EEG data

##### Synthetic data

###### Simulations

To test EPICURUS efficacy and assess generalization across different cortical target, we generated synthetic EEG data across three brain areas: Left hemisphere M1, and DLPFC, and the right IPS. For each target, we generated 100 simulated EEG time-series trials using the FieldTrip toolbox and its SimBio framework [1–3]. EEG data was simulated for a 10-10 electrode configuration (63 channels) placed on a virtual head model based on the MNI152 template. EEG time-series were designed to simulate the temporal and spectral characteristics of typical TMS-evoked responses. The synthetic data were composed of three distinct neural sources: (i) a dipole at the target site (MNI coordinates: Left −M1: −42, −20, 67; Left -DLPFC: −38, 44, 26; or Left -IPS: −40, −46, 46) producing a one-cycle 27Hz oscillation with an average peak-to-peak amplitude of 1µV, starting at a latency of 25ms relative to a virtual TMS pulse. The dipole was oriented towards specific scalp electrodes depending on the targeted area (Left M1: C3; Left DLPFC: F3; Left IPS: P3; 10-10 system); (ii) A dipole in the left somatosensory cortex (Left-S1, MNI coordinates: −48, −21, 50) oriented towards P3 and producing a one-cycle 10Hz oscillation with a peak-to-peak amplitude of 1µV, beginning at 100ms; (iii) Two dipoles the auditory cortical regions, bilaterally (MNI coordinates: −48, −10, −10 and 48, −10, 10) both oriented towards the central midline (Cz), to produce a combined one-cycle 7Hz oscillation with a peak-to-peak amplitude of 2µV, starting at 150ms (Supplementary Figure 1). To simulate realistic variability, small trial-to-trial amplitude variations were introduced (±0.5 µV).

**Supplementary Figure 1.**
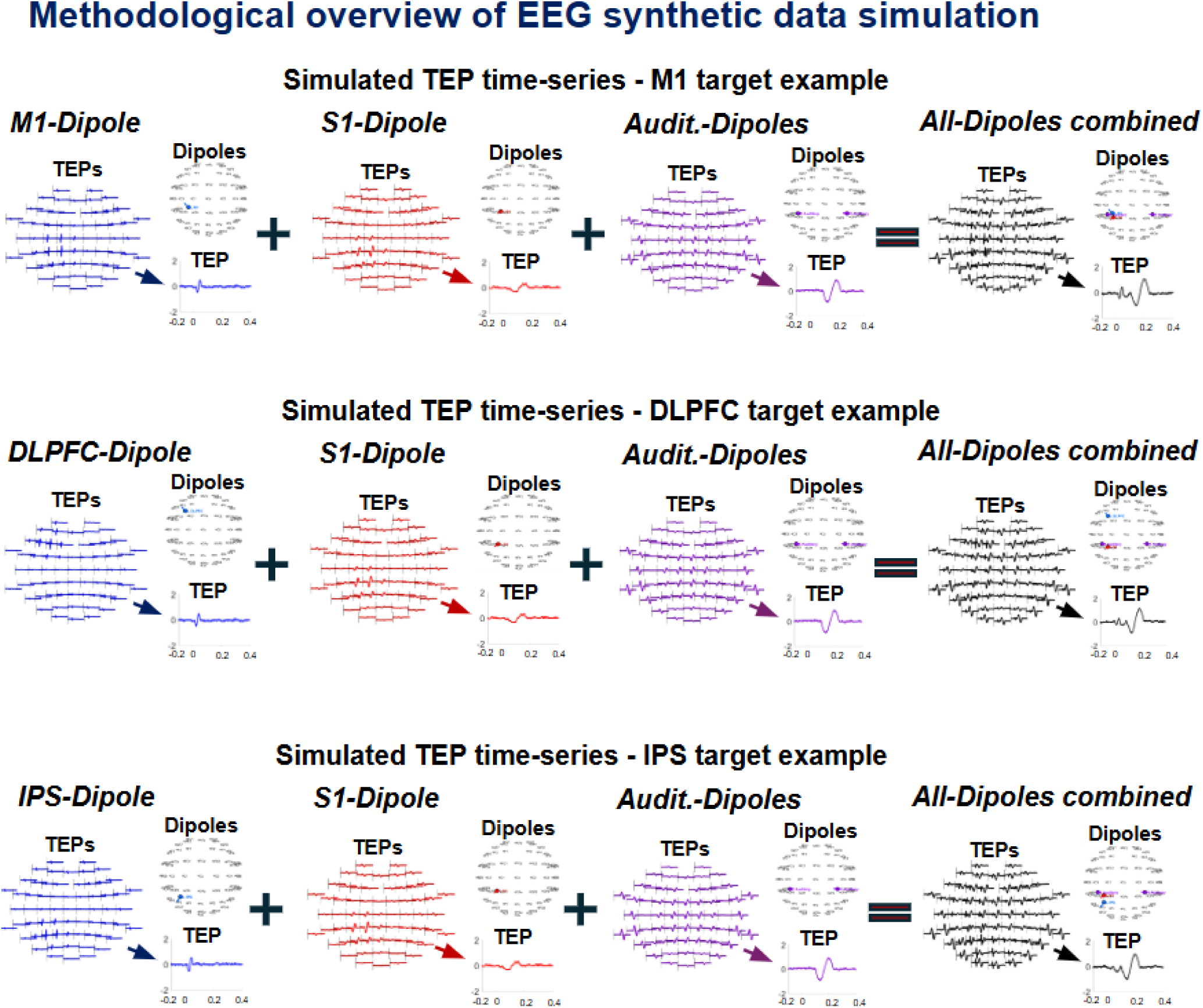
Synthetic data generation. To validate the filter, synthetic datasets were generated for three targets: Primary Motor cortex (M1), Dorsolateral Prefrontal Cortex (DLPFC), and Intraparietal Sulcus (IPS). For each target, independent dipoles were simulated to represent target activity (blue), i.e., the local cortical responses at the stimulation site; the somatosensory indirectly induced activity (red); the auditory indirectly-induced activity (purple). The combined signal (black) represents the ‘synthetic TEP-like response’ used to further compare the extraction efficiency of EPICURUS against prior existing standard methods.

##### TMS-EEG data

###### Participants

To test EPICURUS performance on real human TMS-EEG data, we analyzed a pre-existing dataset consisting of n=18 healthy participants (10 females, 8 males, 26±5 years old) who received TMS over L-M1 while EEG was concurrently recorded. Participants were all right-handed, met the safety criteria for MRI [4] and TMS [5], and provided written consent to participate in the study. Our protocol followed the Declaration of Helsinki and received approval from a local ethical committee (*Comité de Protection des Personnes, Ile de France I*).

**Supplementary Table S1.**
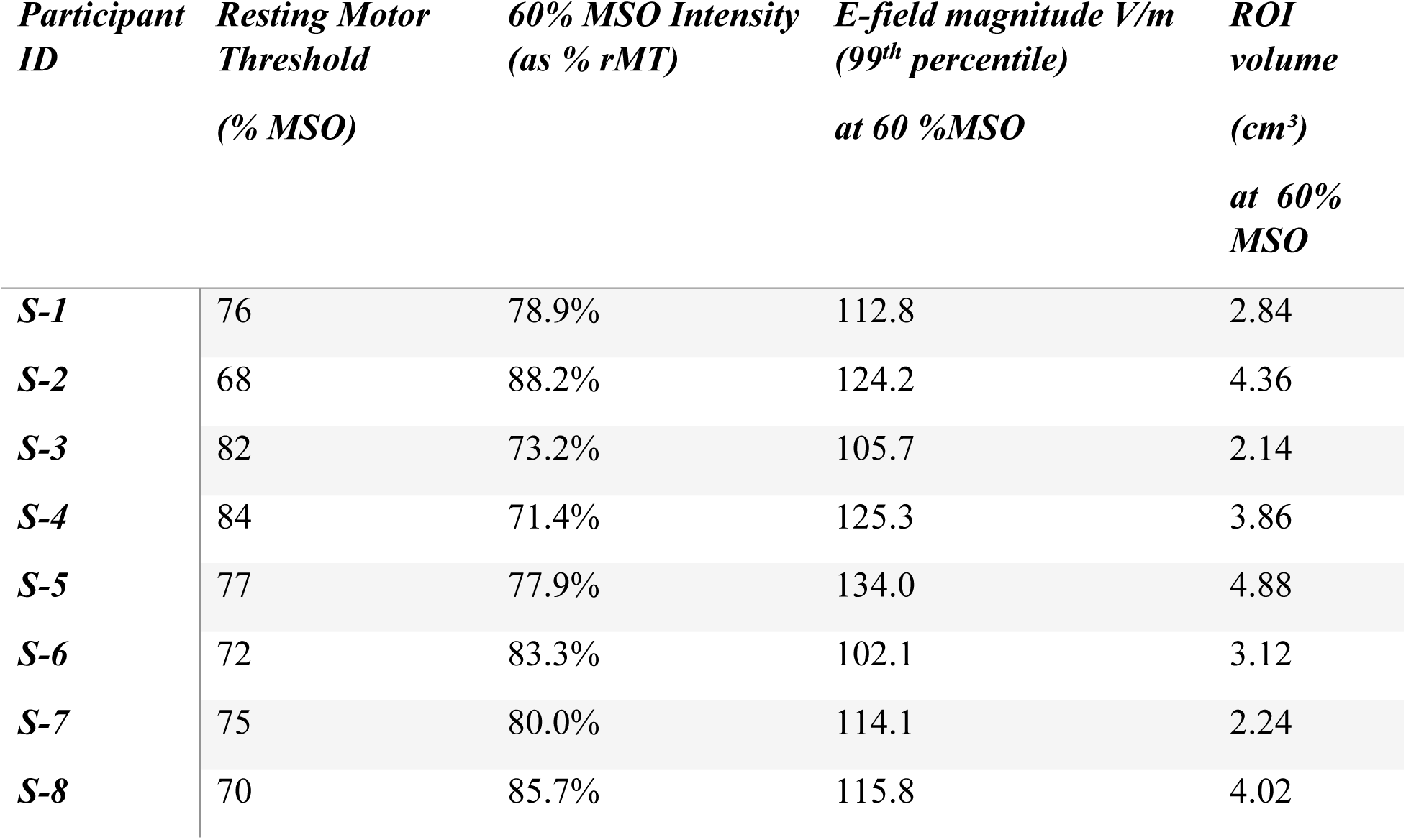
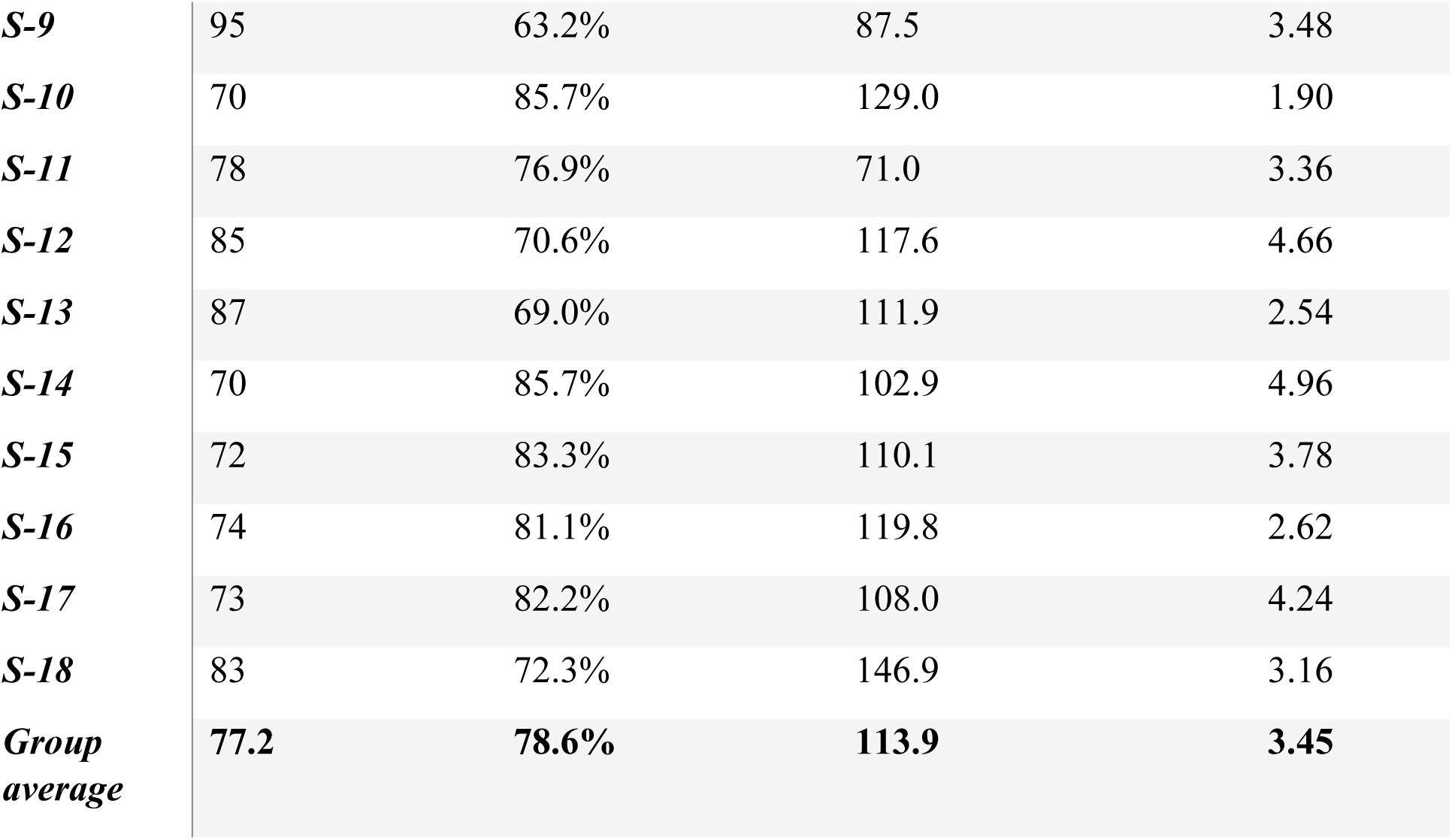
Physiological and biophysical simulation parameters. rMT (% MSO): Resting motor threshold, defined as the minimum TMS intensity required to elicit MEPs > 50μV in 5 out of 10 trials, measured with the EEG cap in place. 60% MSO Intensity (as % rMT): The fixed experimental stimulation intensity (60% MSO) expressed relative to the participant’s individual rMT. E-field magnitude (V/m): The 99th percentile peak electric field magnitude simulated on the gray matter surface at 60% MSO using participant-specific MRI-based models. ROI Volume (cm^3^): The volume of the cortical region-of-interest where the induced electric field exceeded 80% of its maximum magnitude, used to constrain the spatial filter.

###### Data collection

Participants received 80 single TMS pulses (with a variable inter-stimulus interval of 5-7sec) over the left M1, while scalp EEG was continuously recorded (25kHz sampling rate; ActiChamp, with 65 electrodes placed following the International 10/10 system coordinates). TMS was administered with a 70 mm figure-of-eight alpha coil attached to a biphasic stimulator (Magstim Rapid2) at a fixed intensity (60% of the machine stimulation output; MSO; dI/dtmax = 68,8 at 60% MSO [6]). While fixed intensity introduces inter-individual variability in TEPs, prior research suggests that multisensory responses scale more strongly with absolute intensity [7]; thus, a constant intensity would induce comparable multisensory input across participants. Individual resting motor thresholds (rMT), the fixed experimental intensity (60% MSO) expressed as a percentage of the individual rMT, and the simulated electric field magnitudes at 60% MSO are detailed in Table S1. M1 was defined as the optimal cortical location for eliciting consistent MEPs in the First Dorsal Interosseous (FDI) muscle at rest [8]. For each participant, TMS coil positioning was maintained consistently throughout the experiment using an MRI-frameless stereotactic system (Brainsight, Rogue Solutions Inc., Montréal, Québec, Canada), with coordinates and orientations later recorded to inform individualized E-field modeling. To limit TMS-related auditory responses, participants wore foam earplugs through which noise masking was continuously played (ER-3C insert earphones, ETYMOTIC Research) at a volume tailored individually to ensure participants were unable to perceive the TMS clicking sound, or until any further volume increase would be uncomfortable.

###### Data preprocessing

The TMS-EEG data were processed using the MATLAB-based toolbox EEGLAB [9] and its TMS-EEG signal analyzer plugin (TESA [10]). Data were epoched around the TMS pulse (from −1200 to 1200ms -w.r.t. the TMS pulse) and baseline corrected (from −500 to −10ms). TMS-pulse electromagnetic (from −2ms to 12ms w.r.t the TMS pulse) and recharging artifacts (visually inspected [11]) were removed and interpolated using a spline function. The data were down-sampled to 1 kHz, the most artifacted channels were temporarily removed from analyses, and the independent component analysis (ICA) was employed to remove ocular artifacts. Next, the SOUND algorithm was used for noise suppression, replacement of removed channels, and signal re-referencing to a common average [12]. The SSP-SIR algorithm was then used to suppress TMS-evoked muscle artifacts [13], and a second ICA was computed to remove residual continuous muscle artifacts. Finally, the data were bandpass filtered with a fourth-order Butterworth filter (2-45Hz) and baseline corrected.

##### E-field modeling

We modeled the induced E-field for both the synthetic and human data and each stimulation target. For the synthetic data, E-field distributions were modeled using the standardized MNI152 head template provided within the SimNIBS 4.0 environment [14]. For the human data, we used SimNIBS 4.0 and its ‘*charm*’ routine [15] to generate individualized tetrahedral finite element head models from each participant’s structural MRI. E-fields were then estimated by assuming a quasi-static regime with predefined standard isotropic conductivity values. E-field distributions were simulated assuming the use of a D70 mm figure-of-eight alpha coil attached to a biphasic stimulator (Magstim Rapid2) at a fixed intensity (60% MSO; dI/dtmax = 68,8 at 60% MSO [6]). For synthetic data, the coil was centered at the scalp projection closest to the target MNI dipole coordinates, with the handle orientation chosen to reflect standard TMS experimental practice for the corresponding cortical targets. For M1 stimulation, the coil was oriented tangentially to the scalp with the handle pointing posterolaterally at ∼45° to the midsagittal line. For DLPFC and left IPS stimulation, the figure-of-eight coil was held tangentially to the scalp with the handle pointing posteriorly (posterolaterally for DLPFC; posteromedially for IPS), approximately parallel to the midsagittal line. In the case of human data, coil position and orientation were adjusted subject-wise according to the coordinates and orientations recorded during the experiment using a neuronavigation system. From each E-field simulation, the ROI exceeding 80% of the maximum field magnitude (80% to 99.9%) was identified to define the ROI for source-localization of the TMS-EEG signal.

##### Local signal extraction

We extracted the TMS-evoked signals in synthetic (all the target regions) and human data (M1) using four approaches: (1) Sensor-level: the raw signal extracted from the electrode closest to the stimulated target (C3 for M1, F3 for DLPFC, and P3 for IPS); (2) Laplacian: the signal extracted from the same electrode with the average of its immediate neighbors subtracted to increase local contrast; (3) Standard MNE: the mean source signal (Classical L2-MNE based) extracted from the ROI defined by the TMS E-field magnitude (sources exceeding 80% of the E-field; [16]). For MNE source reconstruction, a forward model based on the MNI152 template was first reconstructed. The cortical distributed sources (corresponding to grey matter cortical surface) were formed of freely orienting dipoles using the L2-MNE solution, where source reconstruction was regularized with singular value decomposition using the dimensions corresponding to the 20 largest components [1]; (4) EPICURUS: the source signal extracted using our E-field-based spatial filtering approach optimized using cross-talk functions, incorporating distributed source constraints to minimize signal leakage across the brain and explicit constraints targeting auditory regions to suppress their contributions.

##### EPICURUS: filter design

EPICURUS was built on a previously developed source-space framework (DeFleCT; [17]) that uses crosstalk functions to design and optimize custom spatial filters targeted to a specific ROI. Here, the ROI corresponded to the cortical area targeted by TMS. EPICURUS extended the DeFleCT framework by incorporating the biophysical properties of the TMS-induced E-field, which were used to define the spatial constraints of the filter (Supplementary Figure 2).

The procedure began with the computation of a forward model, derived from a MNI151 template model (for the synthetic data), or from participant-specific MRI (human data) [15,18] to map cortical source activity from sensor-level EEG. The TMS-induced E-field was simulated with SimNIBS and its distribution was used to identify the cortical region most strongly affected by stimulation (i.e., >80% of the maximum E-field magnitude; [16]). For M1 stimulation, defining the core region at this threshold typically captures activity in the gyral crown and lip, which has been shown to be most predictive of motor-evoked potential (MEP) amplitudes. Importantly, this ROI was not treated as a binary inclusion criterion. A custom inverse operator was then derived by enforcing CTF-based constraints within the inversion kernel [18]. These constraints were designed: (i) to progressively attenuate signals as a function of distance, allowing EPICURUS to operate independently of the precise numerical ROI-threshold and to remain robust to variability in the E-field extent; and (ii) to completely suppress signals originating from specific cortical regions (i.e., auditory regions). Auditory regions were bilaterally delineated by registering an anatomical atlas (HCP_MMP1: Human Connectome Project (HCP) Multi-Modal Parcellation) to the head models (using SimNIBS functions) and annotating the mesh elements associated with Brodmann areas 22, 41, and 42 [19].

**Supplementary Figure 2.**
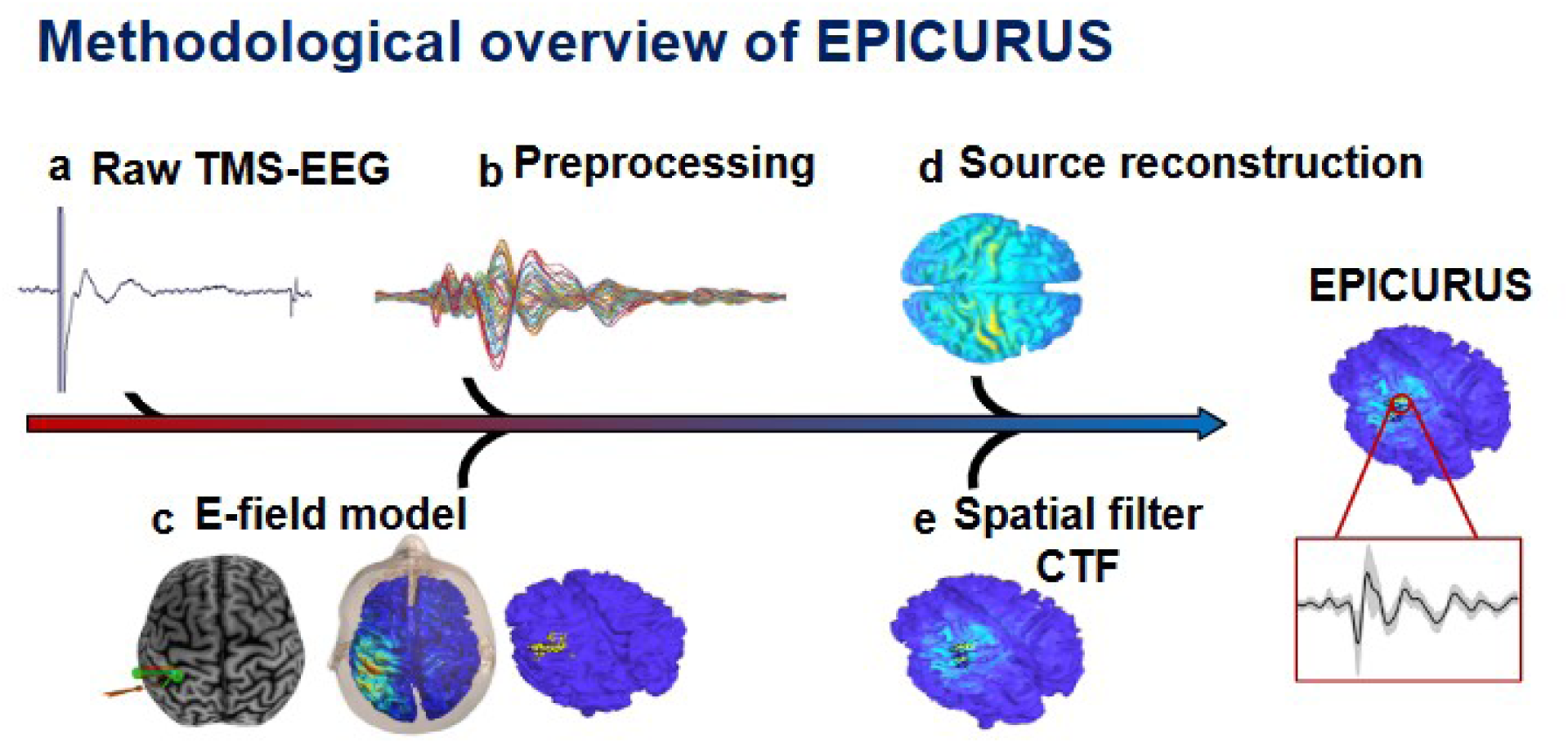
Methodological overview of EPICURUS. The workflow for EPICURUS involves: (**a**) Acquisition of raw TMS-EEG signals; (**b**) Standard TMS-EEG preprocessing; (**c**) Construction of an individualized or template-based E-field model to identify the ROIs corresponding to the TMS stimulated region (i.e., left M1 derived from >80% E-Field strength model); (**d-e**) Source reconstruction, derivation of the EPICURUS spatial filter based on the Cross Talk Functions (CTFs) and extraction of the locally evoked EEG responses via the spatial filter.

#### Statistical analyses

To assess the performance of EPICURUS, TMS-evoked amplitudes and temporal dynamics were compared across the four extraction approaches (Single-sensor, Laplacian, MNE, EPICURUS) for each dataset and target areas, separately. To do this, we used non-parametric, cluster-based permutation statistics (repeated-measures one-way dependent-samples ANOVA; FieldTrip; [2,20]; 2000 permutations; α = 0.05; time window: 0-400 ms relative to the TMS pulse). In the case of simulated data, analyses were performed at the trial level (100 trials per condition, within-subject), whereas in the case of human data, the analysis was performed at the participant level (between-subjects). When significant differences were observed, post-hoc pairwise comparisons between conditions were conducted using non-parametric, independent-samples t-tests (2000 permutations; two-tailed; α = 0.05), with Bonferroni correction applied across pairwise comparisons.

## Extended statistical results

**Supplementary Table 2.** ANOVA statistical results. . Results of the cluster-based permutation tests using a trial-level (in-silico simulated EEG) or group-level (human TMS-EEG) repeated-measures dependent-samples ANOVA.

| Experiment | Target Area | Cluster | Approx. Latency in *ms* | F-statistic | p-value |
| --- | --- | --- | --- | --- | --- |
| In-silico | M1 | 1 | ~20 ms | F(3, 396) = 10.1 | < 0.001 |
|  |  | 2 | ~40 ms | F(3, 396) = 12.3 | < 0.001 |
|  |  | 3 | ~65 ms | F(3, 396) = 14.7 | < 0.001 |
|  |  | 4 | ~145 ms | F(3, 396) = 11.1 | < 0.001 |
|  |  | 5 | ~200 ms | F(3, 396) = 12.8 | < 0.001 |
| In-silico | DLPFC | 1 | ~26 ms | F(3, 396) = 13.9 | < 0.001 |
|  |  | 2 | ~43 ms | F(3, 396) = 14.1 | < 0.001 |
|  |  | 3 | ~70 ms | F(3, 396) = 4.7 | < 0.001 |
|  |  | 4 | ~133 ms | F(3, 396) = 15.9 | < 0.001 |
|  |  | 5 | ~200 ms | F(3, 396) = 5.7 | < 0.001 |
| In-silico | IPS | 1 | ~26 ms | F(3, 396) = 12.8 | < 0.001 |
| | | 2 | ~43 ms | $F(3, 396) = 11.7$ | $< 0.001$ |
| | | 3 | ~80 ms | $F(3, 396) = 5.3$ | $< 0.001$ |
| | | 4 | ~140 ms | $F(3, 396) = 6.5$ | $< 0.001$ |
| | | 5 | ~210 ms | $F(3, 396) = 4.9$ | $< 0.001$ |
| Human | M1 | 1 | ~15 ms | $F(3, 51) = 8.5$ | 0.01 |
| | | 2 | ~40 ms | $F(3, 51) = 7.8$ | 0.002 |
| | | 3 | ~110 ms | $F(3, 51) = 4.3$ | $< 0.001$ |
| | | 4 | ~200 ms | $F(3, 51) = 7.2$ | $< 0.001$ |
| | | 5 | ~325 ms | $F(3, 51) = 6.9$ | $< 0.001$ |

**Supplementary Table 3.** Post-hoc pairwise comparisons results. . Results of the post-hoc tests, non-parametric dependent-samples t-tests (Bonferroni corrected) performed to identify specific differences between pairs of signal extraction strategies.

| Experiment | Target | Comparison | Clusters (Approx. Latency in *ms*) | Key Stats (t-score / p-value) |
| --- | --- | --- | --- | --- |
| In-silico | M1 | Laplacian vs. MNE | 70 | $t = -5.3 / p < 0.001$ |
| | | | 111 | $t = -3.1 / p < 0.001$ |
| | | | 150 | $t = -3.3 / p < 0.001$ |
| | | | 200 | $t = -3.2 / p = 0.001$ |
| | | **Laplacian vs. C3** | 25<br>40<br>135<br>213 | $t = -6.4 / p < 0.001$<br>$t = -5.5 / p < 0.001$<br>$t = -4.9 / p < 0.001$<br>$t = -4.5 / p < 0.001$ |
| | | **Laplacian vs. Epi** | 24<br>40<br>150<br>196 | $t = -6.1 / p < 0.001$<br>$t = -5.9 / p < 0.001$<br>$t = -2.5 / p = 0.02$<br>$t = -3.3 / p = 0.01$ |
| | | **MNE vs. C3** | 22<br>60<br>126<br>210 | $t = -5.6 / p < 0.001$<br>$t = -3.0 / p < 0.001$<br>$t = -5.3 / p < 0.001$<br>$t = -3.1 / p < 0.001$ |
| | | **MNE vs. Epi** | 22<br>76<br>112<br>150<br>205 | $t = -4.6 / p < 0.001$<br>$t = 4.4 / p < 0.001$<br>$t = 2.0 / p < 0.001$<br>$t = 5.0 / p < 0.001$<br>$t = 5.4 / p < 0.001$ |
| | | **C3 vs. Epi** | 69<br>143<br>206 | $t = 2.6 / p < 0.001$<br>$t = 6.2 / p < 0.001$<br>$t = 6.7 / p < 0.001$ |
| **In-silico** | **DLPFC** | **Laplacian vs. MNE** | 70<br>118 | $t = -2.6 / p < 0.001$<br>$t = -2.2 / p < 0.001$ |
| | | **Laplacian vs. F3** | 21<br>45<br>75 | $t = -3.6 / p < 0.001$<br>$t = -5.3 / p < 0.001$ |
| | | | 133<br>200 | $t = -2.5 / p < 0.001$<br>$t = -4.0 / p < 0.001$<br>$t = -2.8 / p < 0.001$ |
| | | Laplacian vs. Epi | 21<br>45 | $t = -4.0 / p < 0.001$<br>$t = -5.7 / p < 0.001$ |
| | | MNE vs. F3 | 22<br>43<br>130<br>200 | $t = -6.1 / p < 0.001$<br>$t = -8.0 / p < 0.001$<br>$t = -2.6 / p < 0.001$<br>$t = -1.9 / p < 0.001$ |
| | | MNE vs. Epi | 22<br>43<br>160 | $t = -6.5 / p < 0.001$<br>$t = -5.9 / p < 0.001$<br>$t = 2.2 / p < 0.001$ |
| | | F3 vs. Epi | 79<br>138<br>206 | $t = 3.9 / p < 0.001$<br>$t = -5.2 / p < 0.001$<br>$t = -3.9 / p < 0.001$ |
| In-silico | IPS | Laplacian vs. P3 | 24<br>40<br>70<br>139<br>206 | $t = 6.1 / p < 0.001$<br>$t = -4.4 / p < 0.001$<br>$t = 3.7 / p < 0.001$<br>$t = 4.6 / p < 0.001$<br>$t = -4.9 / p < 0.001$ |
| | | **Laplacian vs. Epi** | 22<br>40 | $t = 3.6 / p < 0.001$<br>$t = -3.8 / p < 0.001$ |
| | | **MNE vs. P3** | 26<br>40<br>70<br>140<br>205 | $t = 4.5 / p < 0.001$<br>$t = -3.7 / p < 0.001$<br>$t = 3.4 / p < 0.001$<br>$t = 3.7 / p < 0.001$<br>$t = -3.6 / p < 0.001$ |
| | | **MNE vs. Epi** | 22<br>43 | $t = 4.5 / p < 0.001$<br>$t = 2.2 / p < 0.001$ |
| | | **P3 vs. Epi** | 70<br>150<br>206 | $t = -3.9 / p < 0.001$<br>$t = -4.0 / p < 0.001$<br>$t = 3.0 / p < 0.001$ |
| **Human** | **M1** | **Laplacian vs. MNE** | 110<br>200 | $t = -7.3 / p < 0.001$<br>$t = -5.9 / p < 0.001$ |
| | | **Laplacian vs. C3** | 15<br>40<br>115<br>200<br>325 | $t = -4.5 / p = 0.02$<br>$t = -6.7 / p < 0.001$<br>$t = -17.9 / p < 0.001$<br>$t = -15.7 / p < 0.001$<br>$t = -9.5 / p < 0.001$ |
| | | **Laplacian vs. Epi** | 15<br>45<br>95<br>150 | $t = -7.6 / p < 0.001$<br>$t = -2.1 / p = 0.02$<br>$t = -1.9 / p = 0.01$<br>$t = -4.8 / p = 0.001$ |
| | | **MNE vs. C3** | 15<br>38 | $t = 3.3 / p < 0.001$<br>$t = 5.8 / p < 0.001$ |
|  |  |  | **100**<br>**185**<br>**325** | **t = 10.6 / p < 0.001**<br>**t = 9.2 / p < 0.001**<br>**t = 5.4 / p < 0.001** |
|  |  | **MNE vs. Epi** | **14**<br>**110**<br>**185** | **t = -3.3 / p < 0.001**<br>**t = 7.7 / p < 0.001**<br>**t = 4.3 / p < 0.001** |
|  |  | **C3 vs. Epi** | **40**<br>**110**<br>**200**<br>**325** | **t = 1.2 / p = 0.02**<br>**t = -15.2 / p < 0.001**<br>**t = -12.7 / p < 0.001**<br>**t = -8.7 / p < 0.001** |

